# A Systemic and Molecular Study of Subcellular Localization of SARS-CoV-2 Proteins

**DOI:** 10.1101/2020.08.02.233023

**Authors:** Jing Zhang, Ruth Cruz-cosme, Meng-Wei Zhuang, Dongxiao Liu, Yuan Liu, Shaolei Teng, Pei-Hui Wang, Qiyi Tang

## Abstract

Coronavirus possesses the largest RNA genome among all the RNA viruses. Its genome encodes about 29 proteins. Most of the viral proteins are non-structural proteins (NSP) except envelop (E), membrane (M), nucleocapsid (N) and Spike (S) proteins that constitute the viral nucleocapsid, envelop and surface. We have recently cloned all the 29 SARS-CoV-2 genes into vectors for their expressions in mammalian cells except NSP11 that has only 14 amino acids (aa). We are able to express all the 28 cloned SARS-CoV-2 genes in human cells to characterize their subcellular distributions. The proteins of SARS-CoV-2 are mostly cytoplasmic but some are both cytoplasmic and nuclear. Those punctate staining proteins were further investigated by immunofluorescent assay (IFA) using specific antibodies or by co-transfection with an organelle marker-expressing plasmid. As a result, we found that NSP15, ORF6, M and ORF7a are related to Golgi apparatus, and that ORF7b, ORF8 and ORF10 colocalize with endoplasmic reticulum (ER). Interestingly, ORF3a distributes in cell membrane, early endosome, endosome, late endosome and lysosome, which suggests that ORF3a might help the infected virus to usurp endosome and lysosome for viral use. Furthermore, we revealed that NSP13 colocalized with SC35, a protein standing for splicing compartments in the nucleus. Our studies for the first time visualized the subcellular locations of SARS-CoV-2 proteins and might provide novel insights into the viral proteins’ biological functions.

## INTRODUCTION

The family *Corornaviridae* consists of four main subgroups (or genera): α, β, γ, and δ. Αlpha and β genus coronaviruses infect mammals while γ and δ viruses infect primarily birds. So far, only seven coronavirus members from the α and β subfamilies are found to infect humans (1, 2). Alpha- and some β-coronaviruses often infect human but only cause mild diseases such as common cold including HCoV-229E (229E), HCoV-OC43 (OC43), HCoV-NL63 (NL63) and HCoV-HKU1 (HKU1) (1, 3, 4). These 4 coronaviruses contribute to 20% of all the common cold yearly (5), or more than 50% of all the common cold together with Rhinoviruses (6). The patients of common cold present rhinitis and pharyngitis, as well as sneezing, hoarseness, and cough. The illness is usually self-limited and not needed to be hospitalized in healthy people but can be more severe in individuals with heart or lung diseases (6–8). However, some other beta-coronaviruses (CoV) have been imposing tremendous health problem to humans by causing severe acute respiratory syndrome (SARS) (9–11). The current pandemic of beta-coronavirus (SARS-CoV-2) has badly exerted influence on almost all countries, resulting in the disease named COVID-19.

SARS-CoV-2 is a positive-sense single-stranded RNA virus with 29,903 nucleotides that encodes 12 putative open reading frames responsible for about 29 proteins (**NC_045512**). The genes include three groups: a large replicase (rep) gene followed by structural and accessory genes. The first two-thirds of the genome covers the replicase gene that encodes two large ORFs: ORF1a (rep1a) expressing the polyprotein 1a (pp1a) and ORF1b (rep1b) expressing the polyprotein 1ab (pp1ab). Polyproteins, pp1a and pp1ab are then cleaved by viral proteases, PL protease (PLpro) and 3CL protease (3CLpro), also dubbed main protease (Mpro), respectively, into 16 non-structure proteins (NSPs), named from NSP1 through NSP16 (12–14). Polyproteins, pp1a generates the NSP1-11 and pp1ab contains the NSP1–16, respectively. In pp1ab, NSP11 from pp1a becomes NSP12 instead of NSP11 following extension of pp1a into pp1b. Therefore, as shown in figure 1, there are 2 copies of individual proteins of NSP1 through NSP10.

**Figure 1.**
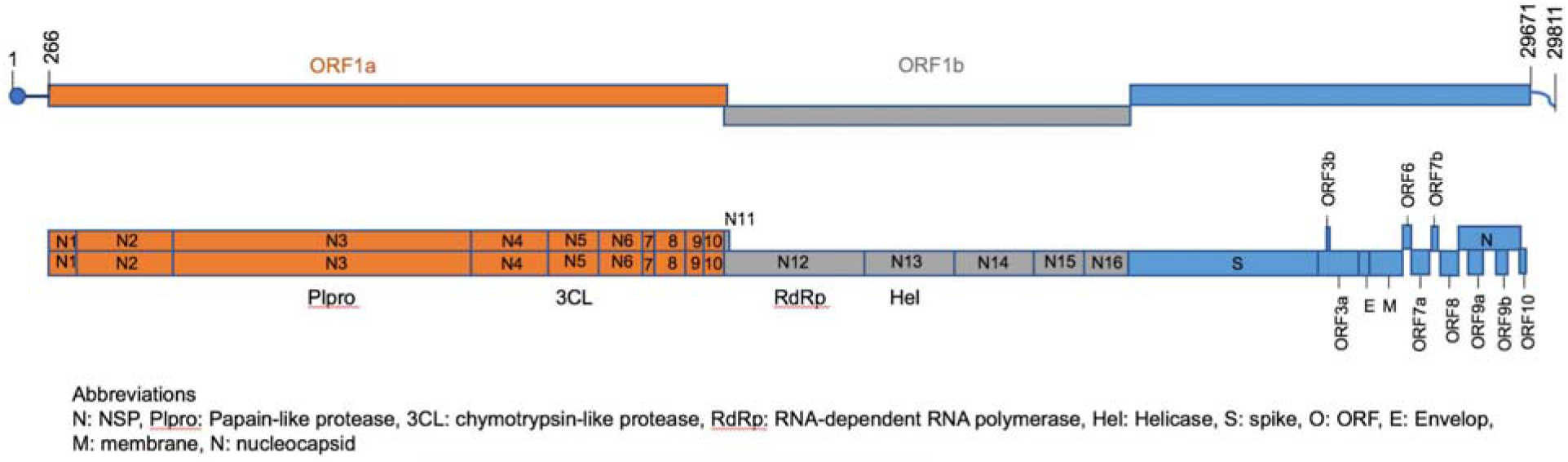
Diagram of SARS-CoV-2 genome and the names of viral genes. The genome of Wuhan-Hu-1 (NC_045512.2) has 29,903 nucleotides and encodes 12 putative open reading frames responsible for about 29 proteins. The first gene, NSP1 starts from 266, and the last gene, ORF10 ends at 29674. The sequence from 1-265 is modified by methylation to form the RNA cap, and from 29672 to 29811 is the 3’ UTR.

Four structural proteins include spike (S), membrane (M), nucleocapsid (N) and envelop (E) proteins. The viral spike protein (S) is cleaved by the human furin enzyme to generate S1, which binds to the host receptor ACE2, and S2, which mediates fusion of virions with the host cell membrane. Both S1 and S2 play crucial roles for viral entry (2). N protein is the only one directly connecting to viral RNA genome. The viral membrane protein (M) is important for maintaining viral structure (15) and the viral envelop protein (E) plays roles in viral assembly and releasing (16, 17). Other accessory proteins encoded by SAR-CoV-2 at its 3’ end are by sub-genomic mRNAs: ORF3a, ORF3b, ORF6, ORF7a, ORF7b, ORF8, ORF9a and ORF9b (some literatures name it as ORF14) and ORF10 (13). The names of the ORFs from ORF8 to the end are not consistent in the current literatures. We adapt the genomic structure and names from https://viralzone.expasy.org/8996 with slight modification: ORF8, ORF9a (was ORF9b), ORF9b (was ORF14) (see figure 1 and table 1).

**Table 1.**
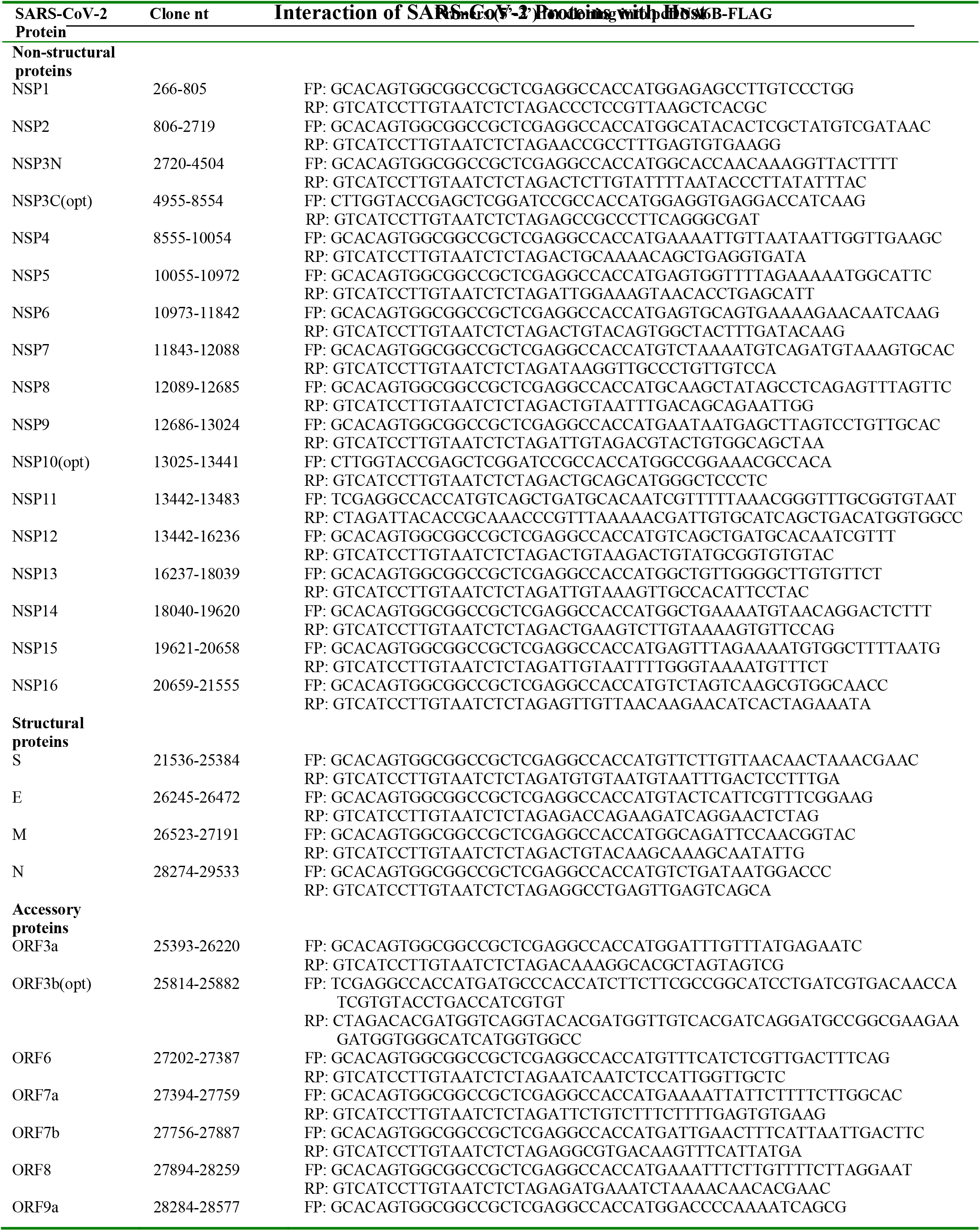
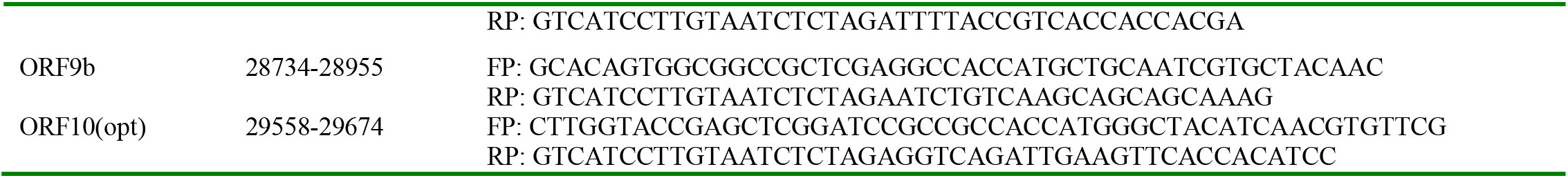
ORFs of SARS-CoV-2 used in plasmid construction. FR: Forward Primer; RP: Reverse Primer; nt: nucleartide; opt: codon optimized.

Each viral protein is potentially important for viral replication by playing a unique biological function. A systemic study of SARS-CoV-2 proteins is still lacking so their functions are largely postulated from SARS-CoV-1 (18). The subcellular distributions of the viral proteins have yet been reported for SARS-CoV-2. It is important to investigate the viral proteins’ locations in cells because the subcellular distribution information not only helps us in understanding how viruses interact with the host cells but also provides clues in fighting against the viral infection. Although related information could be available from SARS-CoV-1, the genomic and protein differences between SARS-CoV-1 and SARS-CoV-2 high likely result in different protein function and subcellular distributions. Therefore, we cloned all the genes of SARS-CoV-2 into vectors for expression in mammalian cells and used immunofluorescent assay (IFA) to examine the viral proteins’ subcellular location in cells. Except for the NSP11 that is only 14 aa long, we are able to express all other 28 viral proteins in Hep-2 cells and found a diversity of protein distribution in cells, suggesting a complicated interaction of SARS-CoV-2 with host cells to achieve a successful infection.

## RESULTS

### 1. SARS-CoV-2 proteins are detected by immunofluorescent assay in human cells

A great number of SARS-CoV-2 strains have been rapidly isolated, and the nucleotide sequences have been accumulating in the GenBank since the COVID-19 pandemic. We selected the Wuhan-Hu-1 (**NC_045512.2**) strain’s sequence to synthesize the DNA fragments for molecular cloning because it was one of the earliest isolates and its sequence has been used for developing the testing and diagnosing methods for COVID-19 (19–21). The DNA fragments of the SARS-CoV-2 genes were synthesized and a FLAG tag was fused in frame to the 3’ terminus of each gene. The genes were cloned into pCAG or pcDNA6B vectors (22, 23). The primers for cloning the genes are listed in table 1. As shown in figure 1, the whole genome contains 29,903 nucleotides (nt). The first viral gene (NSP1) starts from 266 and the last viral gene (ORF10) ends at 29674 (figure 1 and table 1). The exact positions of all genes are listed in table 1. The forward primer (FP) was designated exactly from the starting nucleotide and the reverse primer (RP) was ended at the last nucleotide of each gene. The constructed plasmids were sequenced, and all were confirmed correct. In a systemic attempt of revealing the subcellular locations of SARS-CoV-2 proteins, we transfected each plasmid to HEp-2 cells for 20 hours, then the cells were fixed for immunofluorescent assay (IFA) using anti-Flag antibody to show the viral protein and anti-CoxIV to show the mitochondria or anti-Giantin to show the Golgi apparatus.

The current literatures have been debating regarding the names and numbers of SARS-CoV-2 genes, particularly for the ones after S gene (13, 24). We designed the cloning strategies according to the gene structure published in https://viralzone.expasy.org/8996 with slight modification: 1) ORF3a and ORF3b are the genes after S gene; 2) following the ORF8 are the ORF9a (was ORF9b), ORF9b (was ORF14) and the last one, ORF10 (figure 1). The exact positions of the genes in the viral genome and their cloning primers are listed in table 1. Among the 28 cloned genes of SARS-CoV-2 (NSP-1 through −16 except NSP11, S, M, N, E, ORF3a, ORF3b, ORF6, ORF7a, ORF7b, ORF8, ORF9a, ORF9b and ORF10), their expressions in HEp-2 cells are all detected. A full length NSP3 cloning and expression has not been successful, so we cloned NSP3N and NSP3C both of which can be expressed. The DNA sequences are all correct and the fusion with the FLAG in frame. All the detected SARS-CoV-2 proteins were shown in the figure 1 in green by FITC-tagged anti-FLAG antibody.

As can be seen, the viral proteins are either cytoplasmic (NSP2, NSP3N, NSP3C, NSP4, NSP8, Spike, M, N, E, ORF3a, ORF3b, ORF6, ORF7a, ORF7b, ORF8, ORF9b and ORF10) or both nuclear and cytoplasmic (NSP1, NSP5, NSP6, NSP7, NSP9, NSP10, NSP12, NSP13, NSP14, NSP15 NSP16, and ORF9a). The IFA used a cellular protein antibody against CoxIV to show mitochondria in Texas red-conjugated antibody except those for the NSP9 and M that used anti-Giantin showing Golgi apparatus (red) (figure 2). Although no viral proteins were detected in mitochondria, M protein colocalizes with Giantin, which is consistent to that of SARS-CoV-1 (25, 26). Whether other proteins are related to Golgi apparatus or other cellular organelles needs to be further investigated. Interestingly, some proteins showed punctate staining in the IFA experiments: NSP1, NSP5, NSP9, NSP12, NSP13, NSP14, NSP15, ORF3a and M. The relationships of these proteins with subcellular organelles are further explored in this study.

**Figure 2.**
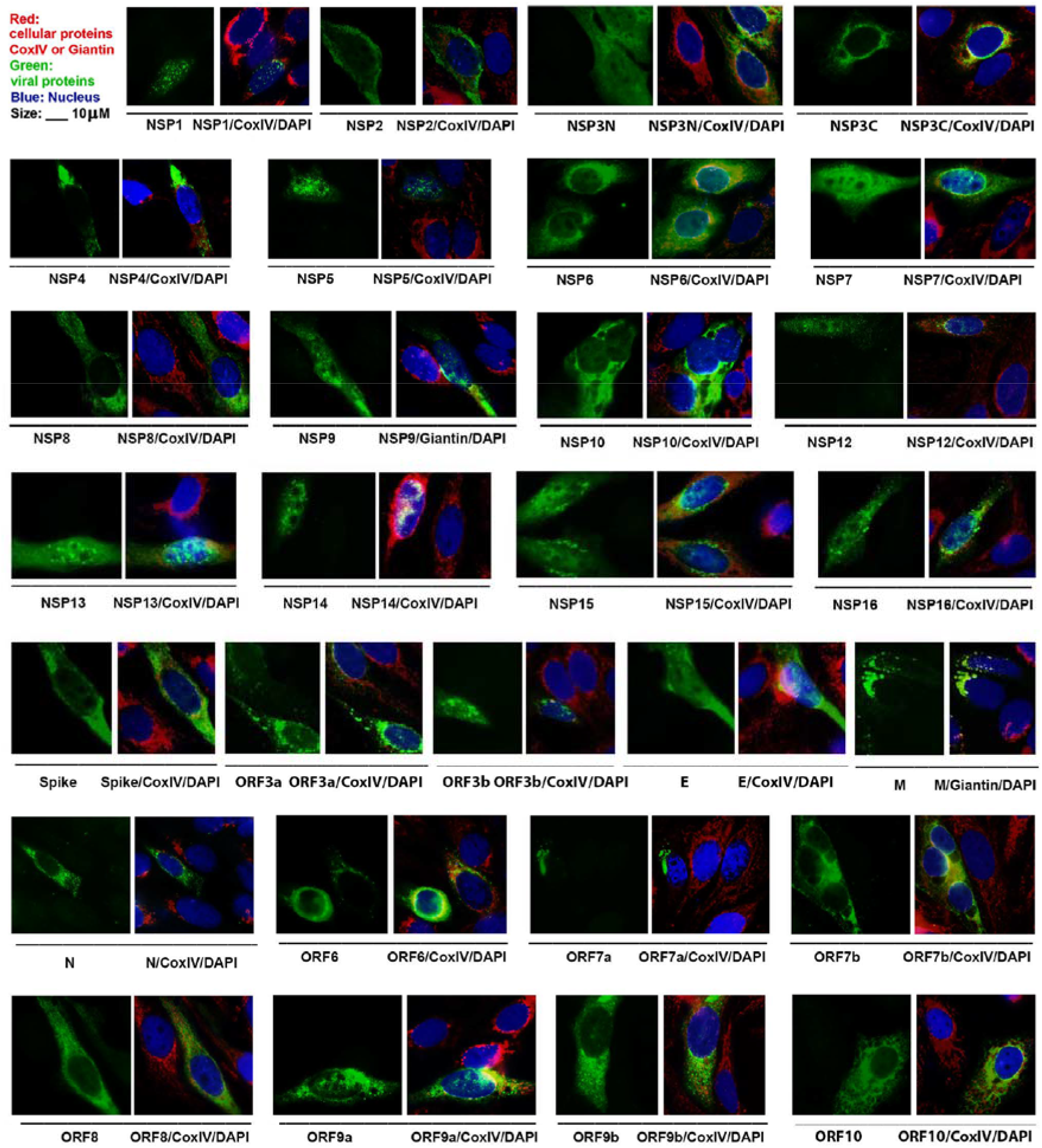
SARS-CoV-2 proteins expressed in human cells. The viral gene fragments (names of the genes are as listed in figure 1 and table 1) were synthesized with a FLAG tag at their 3’ end. Each plasmid was transfected to Hep-2 cells for 20 hours. Immunohistochemistry (IHC) was performed to examine viral protein using anti-FLAG antibody in green (FITC-conjugated 2ndary antibody) and cellular proteins using anti-CoxIV or anti-Giantin antibody in red (Texas red-conjugated 2ndary). The merged pictures include DAPI to show nucleus in blue. Scale bar: 10 μm.

### 2. SARS-CoV-2 proteins, NSP15, M, ORF6 and ORF7a, are associated with Golgi apparatus

Next, we attempted to uncover the subcellular distribution of the viral proteins. Results from figure 2 showed that the proteins encoded by NSP15, NSP16, ORF3, M, ORF6, ORF7a and ORF9a are cytoplasmic punctate proteins. We wondered if they are in any cellular organelles. Here we examined their locations with Golgi apparatus. The Hep-2 cells were fixed at the time of 24 hours post transfection and stained with anti-FLAG in green and anti-Giantin to visualize the Golgi apparatus in red. Consequently, we detected that 4 SARS-CoV-2 proteins are related to Golgi apparatus: NSP15, M, ORF6 and ORF7a. As shown in figure 3, viral proteins M, ORF7a and NSP15 colocalize with Golgi apparatus, and ORF6 partially colocalizes with Golgi apparatus. Except that M-Golgi relationship has been previously reported (26), other proteins’ relationships with Golgi is the first reported by this study.

**Figure 3.**
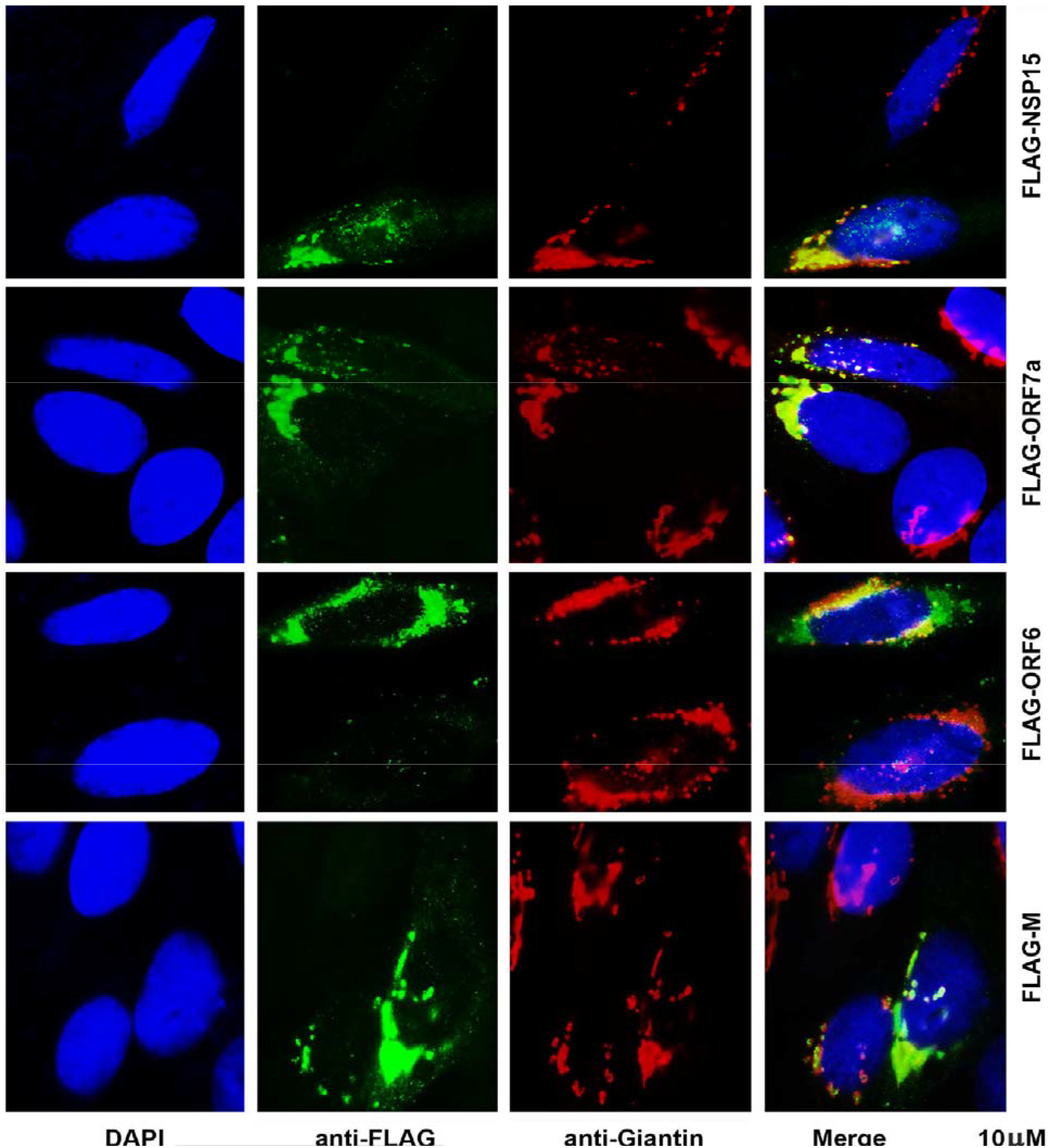
SARS-CoV-2 proteins, NSP15, M, ORF6, and ORF7a, are associated with Golglgi apparatus. The plasmid expressing the protein (indicated on the right side) was transfected to to Hep-2 cells for 24 hours. IFA was performed using anti-FLAG antibody to show viral protein in in green and anti-Giantin antibody to show Golgi apparatus in red. Bar: 10 μm.

A drawback of IFA is that the antibodies could have cross-reaction. To ensure the specificities of the IFA results as shown in figure 3, we employed a co-transfection system using a Golgi protein expression plasmid in which the N-terminus (1-61 Aa) of the Beta-1,4-galactosyltransferase 1 was fused with a cyan fluorescent protein variant, mTurquoise2 (27). In this system, we only need to stain the viral proteins with anti-FLAG antibody. As shown in figure 4, the co-transfected cells were fixed for IFA and the viral proteins were immuno-stained in red fluorescence. DAPI staining is performed to show the nuclei. After merging different colors, the results showed ORF6, ORF7a and NSP15, like M protein, colocalized with Golgi apparatus that was shown by mTurquoise tagged Golgi protein in Cyan. Therefore, we identified 4 SARS-CoV-2 proteins (M, ORF6, ORF7a and NSP15) that are related to Golgi apparatus.

**Figure 4.**
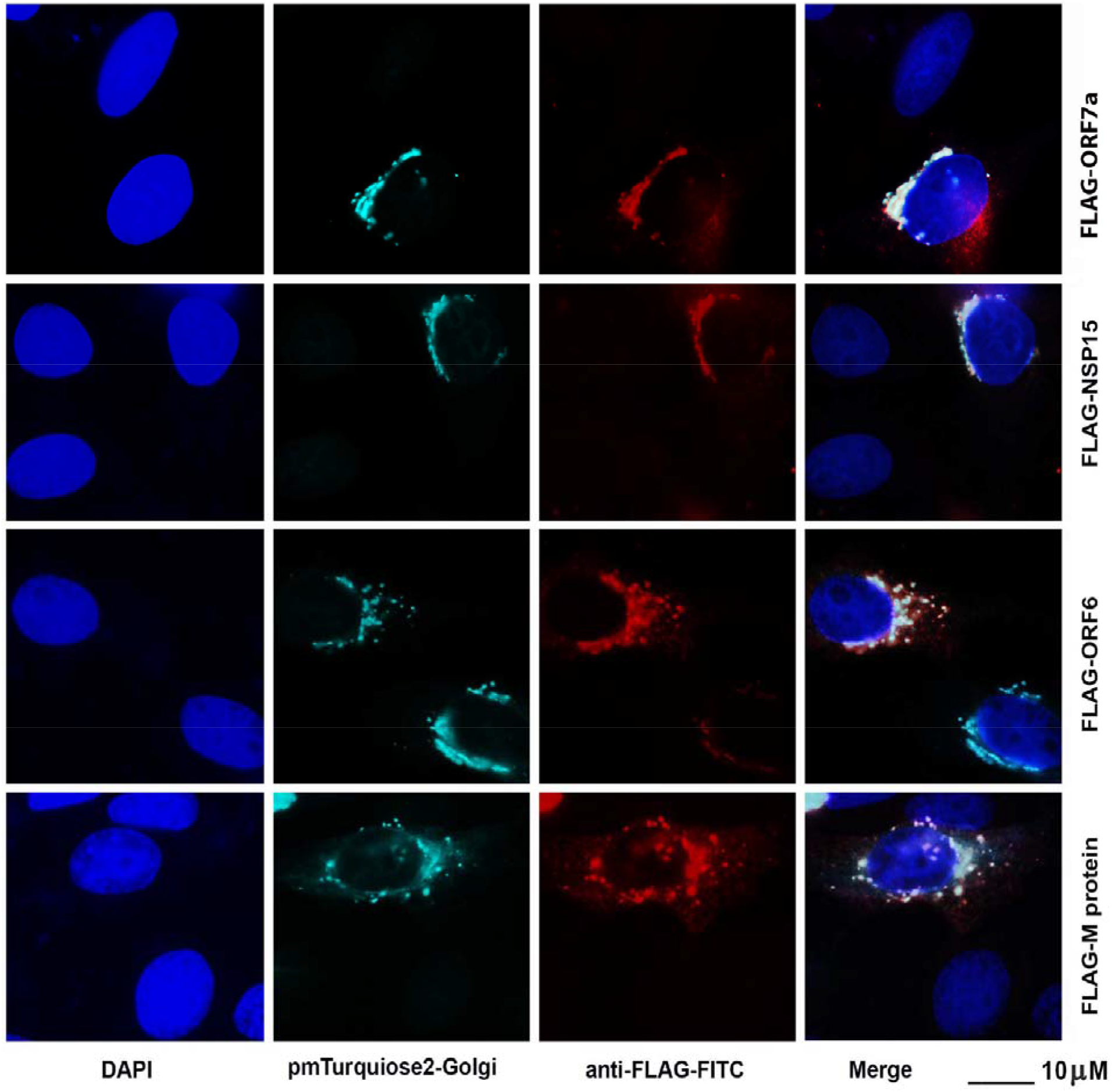
SARS-CoV-2 proteins, NSP15, M, ORF6, and ORF7a, are associated with Golgi apparatus. The plasmid expressing the viral protein (indicated on the right side) was co-transfected with Golgi apparatus marker-expressing plasmid, pmTurquiose2-Golgi, to Hep-2 cells for 24 hours. IFA was performed using anti-FLAG antibody to show viral protein in RED, Golgi apparatus was in Cyan. Bar: 10 μm.

### 3. SARS-CoV-2 protein, ORF7b, ORF8 and ORF10 are related to ER

Like other positive stranded RNA viruses, SARS-CoV-2 RNA is transported to Endoplasmic reticulum (ER) after viral entry. ER is the major cellular organelle that viruses need to usurp because it is a factory for production of viral proteins. Most proteins of SARS-CoV-2 were seen in cytoplasm as shown in figure 2, so we asked whether they colocalize with ER. To that end, we cotransfected several viral protein-expressing plasmids (NSP2, NSP4, NSP6, NSP7, NSP8, NSP9, NSP10) together with pmcCh-sec61-beta (ER and the ER-Golgi intermediate compartment). ER is in red fluorescence because it is tagged with mCherry (28). The viral proteins (ORF7b, ORF8 and ORF10) were shown in green fluorescence by anti-FLAG. Although SARS-CoV-2 proteins are all generated in ER, IFA found only ORF7b, ORF8 and ORF10 colocalized with ER as shown in figure 5. The yellow color in the merged pictures were caused by the colocalization between the viral proteins and Golgi protein: sec61 beta. We also included other proteins (NSP2, NSP4, NSP6, NSP7, NSP8, NSP9, NSP10) in the co-transfection and IFA experiments and they were not colocalizing with ER. ORF7b is a 43 aa protein, ORF8 has only 121 aa and ORF10 contains 38 aa. Although they are small proteins, their functions might be important for viral replication and need to be further investigated.

**Figure 5.**
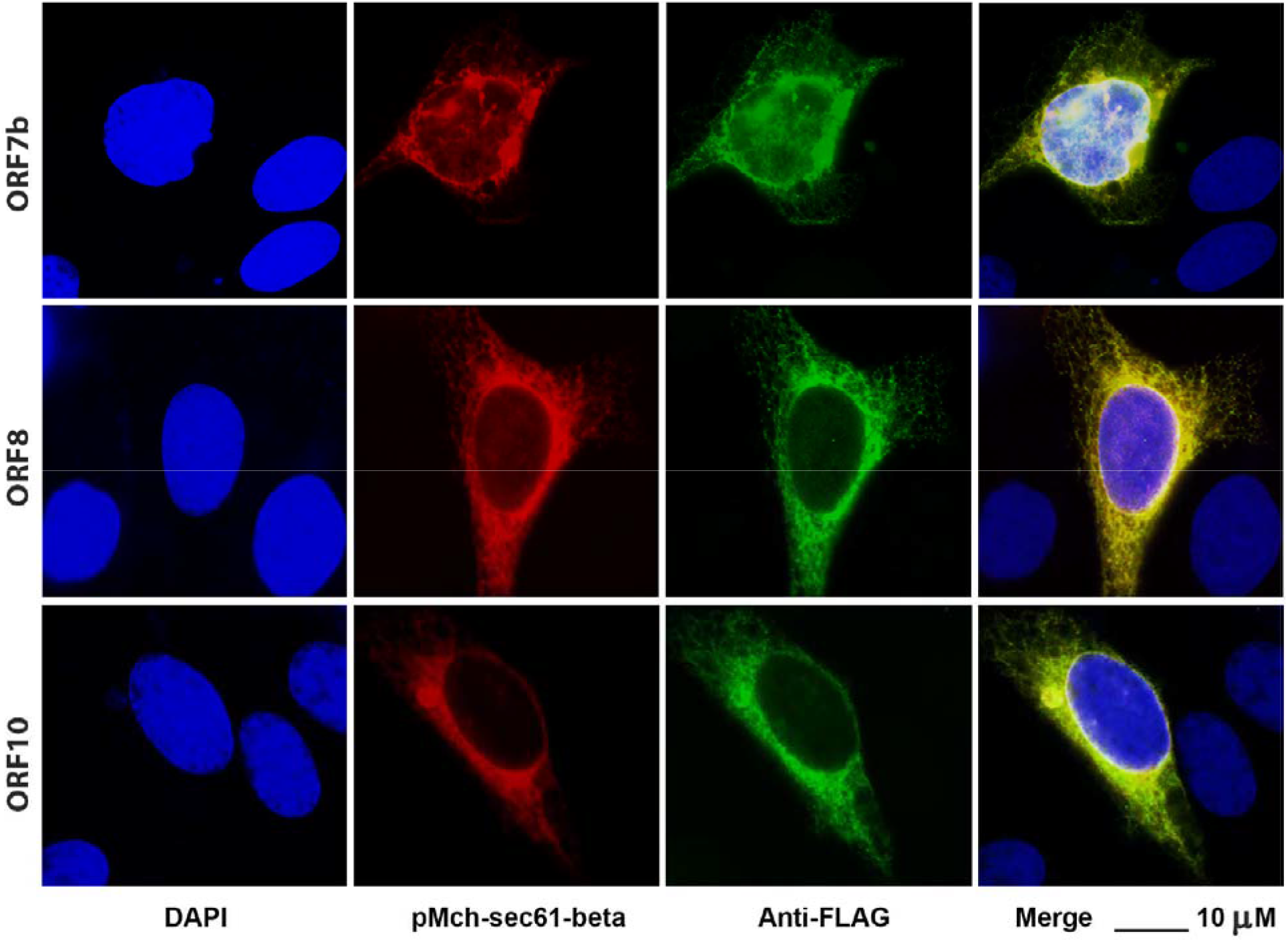
SARS-CoV-2 proteins, ORF7b and ORF10, are related to ER. The plasmid expressing the viral protein (indicated on the right side) was transfected with pMch-sec61-beta to Hep-2 cells for 24 hours. IFA was performed using anti-FLAG antibody to show viral protein in green. ER was shown in red. Bar: 10 μm.

### 4. SARS-CoV-2 protein, ORF3a, is an endosomal and lysosomal protein

The endosomal and the lysosomal systems are important in cellular degradation, reuse of materials, and maintaining homeostasis for cellular living (29). When a molecule is captured through Clathrin-mediated endocytosis, they form the endosome. Endosome is a membrane bound compartment in eukaryotic cells and undergoes a maturation from early endosome to late endosome depending on acidification. The late endosome then fuses with the lysosome to degrade the molecule by lysosomal hydrolytic enzymes. Here we used the plasmids expressing the proteins standing for early endosome, endosome, late endosome, and lysosome, which were cotransfected with SARS-CoV-2 protein-expressing plasmids. We identified ORF3a to be the viral protein that is associated with the formation of endosome and lysosome.

Rab family proteins are usually used as marker of endosomes at different stages: Rab5 is early endosomal protein, Rab11 is an endosomal protein and Rab7 is a late endosomal protein. Lamp1 can be sued to show lysosome (30). Our IFA results showed that only ORF3a is associated with endosome and lysosome. The associations between ORF3a with early endosome, endosome, late endosome and lysosome are shown in figure 6. To confirm the specificity of our IFA assay using the co-transfection system, we also co-transfected ORF3a-expressing plasmid with an ER & Golgi intermediate protein, Rabin8 that is tagged with GFP. No colocalization was detected between ORF3a and Rabin8 (bottom of figure 6). Therefore, we found that ORF3a protein is the SARS-CoV-2 protein related to the endocytosis-related biological activities.

**Figure 6.**
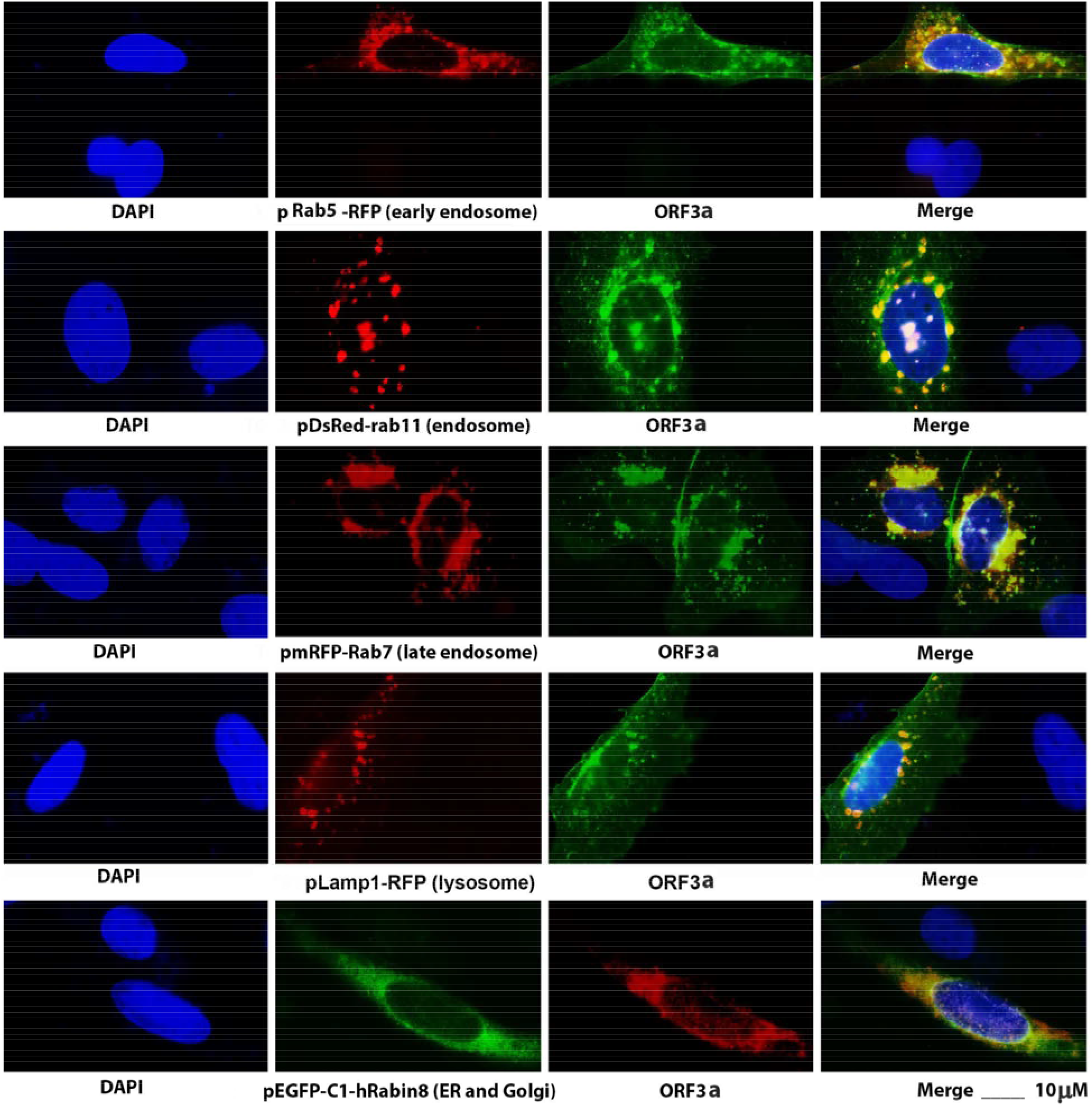
SARS-CoV-2 protein, ORF3a, is related to endosome and lysosome. The plasmid expressing the ORF3a was co-transfected with different cellular marker-expressing plasmid (Rab5, Rab11, Rab7, Lamp1 or Rabin 8) to Hep-2 cells for 24 hours. IFA was performed using anti-FLAG antibody to show viral protein in green or red. Cellular protein was in red or green. Bar: 10 μm.

### 5. SARS-CoV-2 proteins in nucleus

Interestingly, some SARS-CoV-2 proteins are detected in nucleus such as NSP1, NSP5, NSP9 and NSP13 as shown in figure 1. For these nuclear proteins, we decided to know if they are interacting with any nuclear structures such as ND10 (nuclear domain 10 or PML nuclear bodies) that is defensive domain against viral infection or SC (splicing compartment) that is important for gene splicing. To that end, we transfected the NSP1-, NSP5-, NSP9- or NSP13-expressing plasmid to HEp-2 cells for 24 hours. The cells were fixed for IFA with anti-SC35 antibody to show SC or anti-PML antibody to show ND10, and anti-FLAG antibody to show viral proteins. Since no relationship between ND10 and the SARS-CoV-2 proteins were detected, here we only show the IFA results for the SC35.

As shown in figure 7, when we stained the viral proteins, NSP1, NSP5, NSP9 or NSP13 in red and the splicing compartment (SC35) in green, we didn’t detect any relationship between NSP1 and SC35 (bottom of figure 7). Both NSP5 and NSP9 distribute diffusely in the nucleus, but in the strongly stained spots of NSP5 or NSP9, SC35 appears to colocalize with the viral proteins (middle 2 panels of figure 7). Interestingly, NSP13 exists in the nucleus as round “dots” that exactly colocalize with SC35 as can be seen in the top panel of figure 7. This is very interesting because the SARS-CoV-2 is an RNA virus and thought not to involve any gene-splicing activity in the nucleus. This phenomenon was also found for Zika virus that produce NS5 to interact with SC35 (31). However, the consequence of the interaction between RNA viruses and SC35 remains enigmatic and is important to be further investigated.

**Figure 7.**
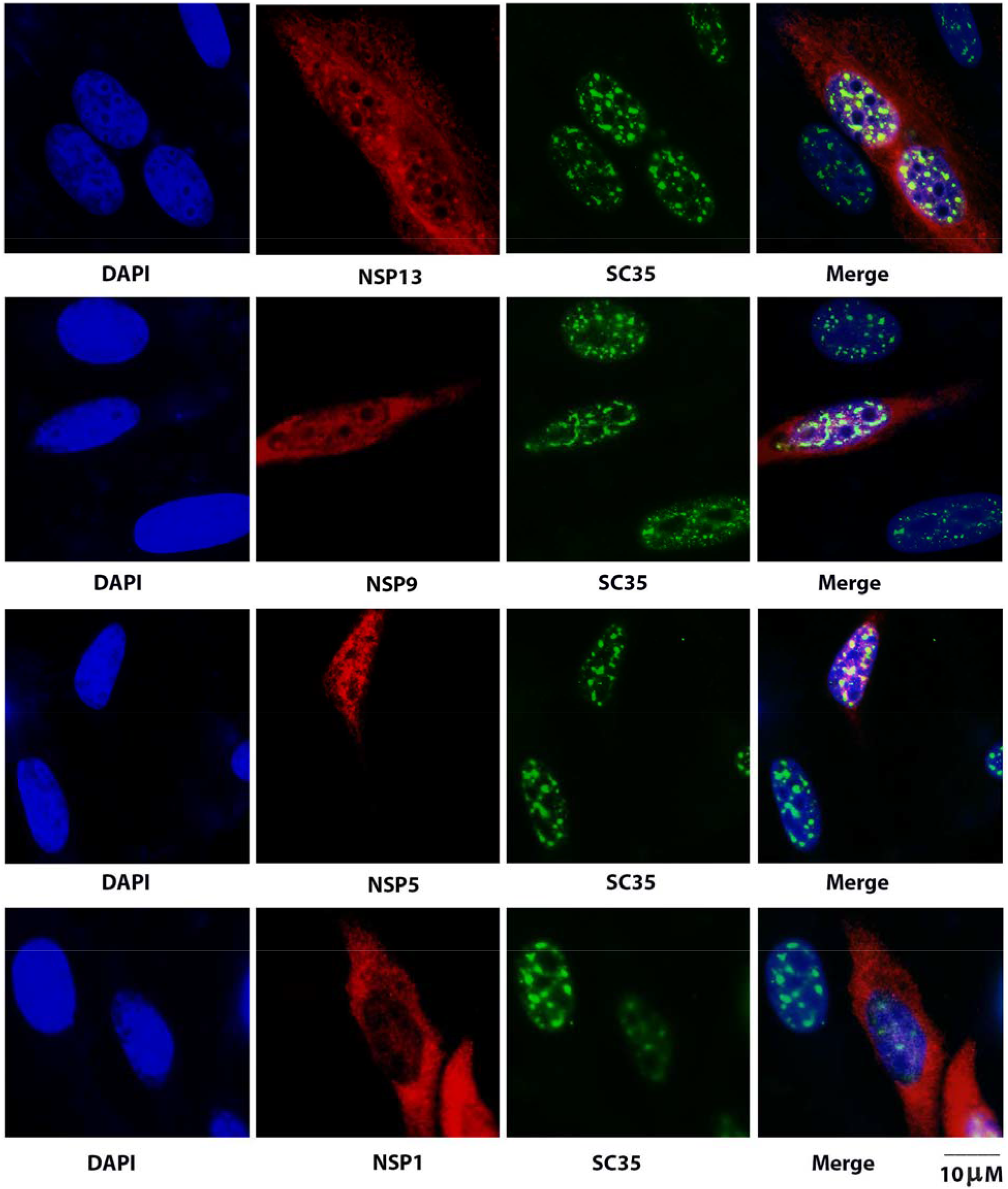
SARS-CoV-2 proteins in nucleus. The plasmid expressing the viral protein as indicated was transfected to Hep-2 cells for 24 hours. IFA was performed using anti-FLAG antibody to show viral protein in red and anti-SC35 antibody to show SC in green. Bar: 10 μm.

## DISCUSSION

The major contribution of this study to the field of COVID-19 is that we have systemically investigated the subcellular distribution of SRAS-CoV-2 proteins. To our knowledge, it is for the first time to clone all the genes of SARS-CoV-2 except NSP11 that is only 14 aa long. In addition, full length NSP3 was cloned but failed to be expressed, so we separated NSP3 into NSP3N and NSP3C. Our overarching findings include that most SARS-CoV-2 proteins are cytoplasmic, and some are both cytoplasmic and nuclear proteins, that 4 proteins in total are detected in Golgi apparatus, that ORF3a colocalizes with endosome and lysosome, and that ORF7b, ORF8 and ORF10 colocalize with ER. Although M protein in Golgi apparatus was previously reported for SARS-CoV-1 (25, 26), it is interesting that M protein is not the only one associated with Golgi apparatus, suggesting that M protein might not be sufficient to ensure the maturation of E and S proteins, some non-structural proteins such as NSP15, ORF6 and ORF7a might be needed for viral maturation. The novel information might implicate biological functions for the viral proteins and help developing new anti-viral strategies against COVID-19.

The biological activities in Golgi apparatus is extremely important for coronaviruses because they assemble and maturate their particle in Golgi apparatus, which differs from most enveloped RNA viruses whose particles assemble and mature in cell membrane (32–35). Coronavirus surface consists M, E and S proteins and they exist differentially in cells. It was detected that S is in several subcellular compartments along the secretory pathway from the ER to the plasma membrane, that M predominantly localizes in the Golgi, where it accumulates, and in trafficking vesicles, and that the E protein is found in perinuclear patches colocalizing with ER (25). To have an efficient virion assembly, the M, E and S proteins must be targeted to the budding site (ER and Golgi apparatus) and to interact with each other or the ribonucleoprotein (N). Therefore, the efficient incorporation of viral E, S and M proteins into CoV virions depends on protein trafficking and protein–protein interactions near the ERGIC (ER Golgi intermediate compartment). It was thought that M protein plays role in the interaction among the viral structural proteins (26, 36). Here we found more coronavirus proteins in Golgi apparatus, especially ORF6, ORF7a and NSP15 (figures 3 and 4). In addition, we found ORF7b, ORF8 and ORF10 colocalize with ER (figure 5). E protein was reported to be in ER for SARS-CoV-1 (26), but not for SARS-CoV-2 in this study. Therefore, we postulate that ORF6, ORF7a, ORF7b, ORF8, ORF10 and NSP15 might also be important in assembly and maturation of SARS-CoV-2.

Endocytosis and endosome acidification inhibitors inhibit infection by CHIKV, murine leukemia virus (MLV), or SARS-CoV-1, implicating that these viral entries into host cells happen through endosomes and require endosome acidification (37). It is known that SARS-CoV-2 depends on the S protein to enter permissive cells. However, it is unknown whether other proteins of SARS-CoV-2 is needed for the entered virus to be successfully trafficked into ER for viral replication and protein synthesis. Here, we found that a non-structural protein, ORF3a, is interacting with all stages of endosome/lysosome system (figure 6). ORF3a colocalizes with early endosome protein Rab5, recycling endosome protein Rab11, late early some protein Rab7 and lysosome protein Lamp1, suggesting the ORF3a protein participated in the endocytic procedure that finally leads the fusion of endosome and lysosome. Whether or not the ORF3a involves in the cellular self-clearance of the toxic materials so that the infected SARS-CoV-2 replicate better remains to be investigated.

Most RNA viruses replicate RNA, translate protein and assemble viral particles in cytoplasm, so the results that some viral proteins colocalize with SC35 (figure 7) are surprising. SARS-CoV-2 RNA doesn’t need a gene splicing. At the beginning to characterize the subnuclear distribution of these potential nuclear proteins (NSP1, NSP5, NSP9 and NSP13), we tested other nuclear domains such as ND10. These viral proteins are not related to ND10 (data not shown). NSP13 forms “dot” like structure in the nucleus and colocalizes with SC35 that is usually used as the marker of splicing compartment (SC). We previously also detected that Zika virus (ZIKV) NS5 not only physically interacts with SC35 in the nucleus by IFA but also molecularly interacts with SC35 by co-immunoprecipitation assay (31). Since this happens often to RNA viruses, it is important for us to ascertain whether SC (or just SC35) potentially involve in viral gene expression or viral replication.

In summary, we molecularly cloned all the genes of SARS-CoV-2 and applied a systemic IFA to characterize the subcellular distribution of the viral proteins. Except NSP11 that were not expressed for unknown reason, we figured out their distribution of total 28 proteins in cells. More importantly, we revealed that more proteins (M, NSP15, ORF7a and ORF6) are related to Golgi apparatus, that ORF7b, ORF8 and ORF10 interacted with ER, that ORF3a participated in all the stages of endosome/lysosome system, and that NSP13 colocalized with SC35, a gene-splicing protein in the nucleus.

The drawback of our study is that we used cloned plasmids that only expresses a viral protein at one time. Transfection system could be different from the viral infection system. Therefore, a detailed study should be conducted in the context of infection of SARS-CoV-2 in cells. In addition, colocalization of the viral proteins with the cellular organelle doesn’t completely stand for the interaction of two proteins, a co-IP is needed to confirm the protein-protein interaction. It is especially important for ORF3a because its function apparently relates to cell membrane, endosome, lysosome and probably phagosome. However, due to the biosafety restriction and lack of a whole set antibodies to the SARS-CoV-2 proteins, our transfection system using the cloned plasmids provides a first-hand insight into the interaction of viral proteins and host cells.

## MATERIALS AND METHODS

### Molecular cloning

The sequence of severe acute respiratory syndrome coronavirus 2 (SARS-CoV-2, **GenBank: NC_045512.2**) isolate, Wuhan-Hu-1, was used in this study for DNA synthesis of each gene (General Bio, China). Among them, the DNA sequences of NSP3C, NSP10, ORF3b, and ORF10 were codon optimized to get a high expression level in human cells. The viral genes were firstly cloned into pcDNA6B-FLAG vector using the Seamless Cloning Kit (Beyotime Biotechnology, China) or standard molecular cloning methods as previously described (38). To obtain a high expression level, these viral genes were then subcloned into pCAG-FLAG. The information about all SARS-CoV-2 genes was listed in table 1.

### Cell lines and Tissue culture

Hep-2 cells (ATCC® CCL-23™) were purchased from ATCC. The cells were maintained in Dulbecco’s modified Eagle’s medium (DMEM) supplemented with 10% fetal calf serum (FCS) and penicillin (100 IU/ml)-streptomycin (100 ug/ml) and amphotericin B (2.5 ug/ml) (39).

### Plasmids and transfection

To show a protein’s location in the Hep-2 cells, the plasmid constructed from “molecular cloning” is co-transfected with one of the following plasmids that expresses a cell marker. pmTurquiose2-Golgi (Beta-1,4-galactosyltransferase 1 1-61 Aa) in red is to show the Golgi apparatus (Addgene cat 36205) (27); pMch-sec61-beta shows ER and the ER-Golgi intermediate compartment (Addgene cat 49155) (40); pLamp1-RFP represents lysosome (Addgene cat 1817) (41); pDsRed-rab11 visualizes endosome (42); pmRFP-Rab7 shows late endosome (42); pmRFP-Rab5 shows early endosome (42); and pEGFP-C1-hRabin8 shows the intermediate cisternae between ER and Golgi (42). These plasmids are purchased from Addgene (http://www.addgene.org). Transfection reagent, Lipofectamine 3000, was purchased from Invitrogen and used according to the manufacturer’s protocol.

### Antibodies

Anti-Giantin (ab80864) for visualizing Golgi body, and anti-CoxIV (ab16056) for showing mitochondria were purchased from Abcam (Cambridge, MA). Anti-Tubulin (4G1, sc-58666) were purchased from Santa Cruz Biotechnology (Santa Cruz, CA). Anti-FLAG antibody (monoclonal), M2, and the anti-SC35 antibody (S4045) were purchased from Sigma.

### Immunofluorescent assay (IFA)

Immunostaining was performed on cells grown on coverslips after fixation with 1% paraformaldehyde (10 min at room temperature) and permeabilization in 0.2% Triton (20 min on ice) by sequential incubation with primary and Texas red (TR)-labeled secondary antibodies (Vector Laboratories, Burlingame, Calif.) for 30 min each (all solutions in PBS). Finally, cells were equilibrated in PBS, stained for DNA with Hoechst 33258 (0.5 μg/ml), and mounted in Fluoromount G (Fisher Scientific, Newark, Del.).

### Confocal microscopy

Cells were examined with a Leica TCS SPII confocal laser scanning system. Two or three channels were recorded simultaneously and/or sequentially and controlled for possible breakthrough between the fluorescein isothiocyanate and Texas Red signals and between the blue and red channels.

## ACKNOWLEDGEMENTS

This study was supported by an NIH/NIAID SC1AI112785 (Q.T.), an NIH/DE R01DE028583-01 (subaward to Q.T.), and National Institute on Minority Health and Health Disparities of the National Institutes of Health under Award Number G12MD007597. This work was supported by grants from the COVID-19 emergency tackling research project of Shandong University (Grant No. 2020XGB03 to P.-H.W) and grants from the Natural Science Foundation of Jiangsu Province (SBK2020042706 to P.-H.W) We thank Translational Medicine Core Facility of Shandong University for consultation and instrument availability that supported this work.

